# Recurrent neural networks that learn multi-step visual routines with reinforcement learning

**DOI:** 10.1101/2023.07.03.547198

**Authors:** Sami Mollard, Catherine Wacongne, Sander M. Bohte, Pieter R. Roelfsema

## Abstract

Many cognitive problems can be decomposed into series of subproblems that are solved sequentially by the brain. When subproblems are solved, relevant intermediate results need to be stored by neurons and propagated to the next subproblem, until the overarching goal has been completed. We will here consider visual tasks, which can be decomposed into sequences of elemental visual operations. Experimental evidence suggests that intermediate results of the elemental operations are stored in working memory as an enhancement of neural activity in the visual cortex. The focus of enhanced activity is then available for subsequent operations to act upon. The main question at stake is how the elemental operations and their sequencing can emerge in neural networks that are trained with only rewards, in a reinforcement learning setting.

We here propose a new recurrent neural network architecture that can learn composite visual tasks that require the application of successive elemental operations. Specifically, we selected three tasks for which electrophysiological recordings of monkeys’ visual cortex are available. To train the networks, we used RELEARNN, a biologically plausible four-factor Hebbian learning rule, which is local both in time and space. We report that networks learn elemental operations, such as contour grouping and visual search, and execute sequences of operations, solely based on the characteristics of the visual stimuli and the reward structure of a task.

After training was completed, the activity of the units of the neural network elicited by behaviorally relevant image items was stronger than that elicited by irrelevant ones, just as has been observed in the visual cortex of monkeys solving the same tasks. Relevant information that needed to be exchanged between subroutines was maintained as a focus of enhanced activity and passed on to the subsequent subroutines. Our results demonstrate how a biologically plausible learning rule can train a recurrent neural network on multistep visual tasks.

**Author Summary:** Many visual problems, like finding your way on a map, are solved by decomposing them into a series of subproblems. For a successful decomposition, the neuronal processes that solve one subproblem must make their results available to the subsequent ones. Experiments in monkeys demonstrated that the outcomes of subproblems are represented as foci of enhanced activity in the visual cortex, which are related to attention shifts and can be used as inputs for the processes solving the next subproblems. To understand how subproblems and their sequencing are learned, we trained a recurrent artificial neural network on the same tasks that monkeys performed, using a biologically plausible reinforcement learning rule. The networks learned the tasks and, importantly, the activation of the networks’ units resembled the spatiotemporal patterns of activity observed in the visual cortex of monkeys. Our results shed light on how recurrent neural networks trained with a biologically plausible learning rule can learn to propagate enhanced activity between subroutines to solve complex visual tasks.

## Introduction

Many everyday tasks can be decomposed into subroutines that are executed sequentially. If you want to find your way on a map, for instance, you first search where your goal and actual position are, and then you find the path between those two positions. Ullman [1] proposed that such complex visual tasks can be solved using visual routines, neuronal programs consisting of a sequence of atomic mental steps that he called elemental operations. Curve tracing, visual search, cuing, matching or contour grouping are examples of such elemental operations [2–4]. The visual routine hypothesis has been applied in various domains, including visual search [5], map reading [3], driving [6,7], block copying [8,9] and other visual tasks [10].

There are at least two critical conditions for the successful implementation of visual routines: the correct elemental operations must be chosen and applied in the appropriate order, and information must flow between subroutines so that the output of one subroutine can be used as the input for the next one. Those two conditions are straightforward to implement in a computer program, but it remains unclear how variables are bound to subroutines in the brain. Several studies provided insights into the neural basis of elemental operations and visual routines. For example, Jolicoeur et al. [5] focused on curve tracing as an example elemental operation. In their task, subjects had to determine if two dots were on the same or different curves. They found that curve-tracing is a serial mental operation because the reaction time increases when subjects trace longer curves (see Fig. 1A for an example curve-tracing stimulus).

**Fig. 1.**
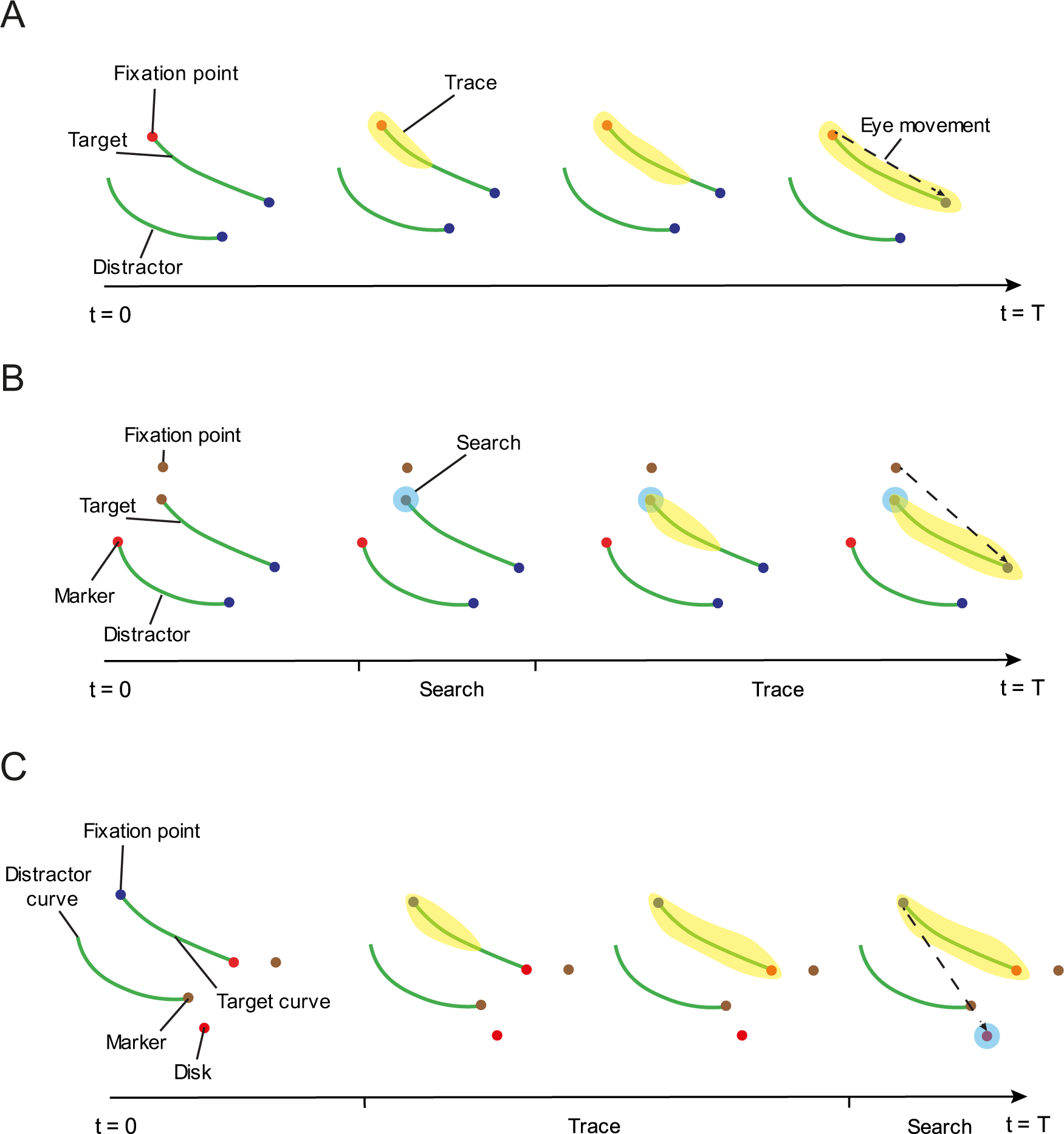
Example stimuli for the three tasks. **A,** Trace task. The monkey makes an eye movement toward the blue dot connected to the red fixation point. The representation of the target curve in the visual cortex is enhanced because extra neuronal activity spreads over this curve (yellow). **B,** Search-then-trace task. The monkey searches for a marker with one of two colors and then traces the curve that starts at this marker to its other end to make an eye movement to the blue dot. This task requires visual search followed by curve-tracing. In the visual cortex, the search operation first labels the target marker with enhanced activity (light blue circle). Its position can be used as the starting position of the tracing operation which propagates enhanced activity over the target curve (light yellow)**. C,** Trace-then-search task. The monkey first traces the target curve connected to the fixation point and identifies the color at the end of this curve. It then has to search for a disk with the same color, which is the target for an eye movement. In the visual cortex, enhanced activity first propagates across the target curve and identifies the target color (trace operation, light yellow), which is used during the subsequent search (light blue circle).

Previous studies started to gain insight in how elemental operations are implemented in the visual cortex. The neural activity during elemental operations can be divided in several phases [12,13]. The first phase is dominated by feedforward processing. Basic features are registered in low level visual areas and more complex features, such as object category are registered in higher areas [14,15]. Later phases are dominated by recurrent processing, enabled by feedback and horizontal connections, allowing the early visual cortex to act as a cognitive blackboard [16]. Higher areas may write to the blackboard in early visual cortex using feedback connections, to enhance the representation of task-relevant features and suppress the representation of distracting information. Other higher visual areas can then read from the blackboard utilizing feedforward connections to selectively process features that are task relevant.

Studies in monkeys carrying out attention-demanding tasks have supported the blackboard analogy. For example, Roelfsema et al. [11] trained monkeys in a curve-tracing task while recording the activity of neurons in the primary visual cortex (area V1). The animals were trained to make an eye movement toward the end of a target curve that was connected to the fixation point, while ignoring another distractor curve (Fig. 1A). The initial response of the neurons was determined by the information in the neurons’ receptive field that is carried by feedforward connections from the retina. However, at a later phase of the neuronal responses, neurons with a receptive field (RF) on the target curve enhanced their activity compared to neurons whose RF fell on distractor curves. This later response modulation reflects an influence of recurrent processing by horizontal connections within V1 and feedback from higher visual areas [15]. Later studies demonstrated that the enhanced activity spreads gradually over the V1 representation of the curve [18]. The latency of the response enhancement was short at the start of the curve, and it occurred later for neurons with RFs farther along the curve. These findings demonstrate the serial nature of the contour-grouping process. Related findings have been reported for visual search, during which the representation of items that are searched for is enhanced during a delayed phase of the neuronal responses in the monkey visual cortex [19–24]. The early feedforward processing phase and the later recurrent processing phase map onto what Ullman called the base- and incremental representation [1,15].

To examine how elemental operations can be combined into visual routines, Moro et al. [19] and Roelfsema et al. [20] recorded from neurons in the visual cortex of monkeys trained to perform more complex visual tasks. The first task was a ‘search-then-trace’ task (Fig. 1B). In this task, the fixation point could be one of two colors. The monkey had to register the color of the fixation point and to search for a circular marker with the same color, which demarcated the start of the target curve. The monkey then traced the target curve to its other end and planned an eye movement to a circle at that position (blue dot in Fig. 1B). The search and trace operations were performed mentally, while the monkey kept its gaze on the fixation point. The eye movement was made only at the end of the trial and the monkey received a reward when he made an eye movement to the circle at the end of the target curve. The second composite task was a trace-then-search task (Fig. 1C). Now the order of the two subroutines was reversed: the monkey first had to trace the target curve connected to the fixation point and to register the color of a marker at the end of this curve. The monkey then searched for a larger circle with the same color that was the target for an eye movement.

The neurophysiological studies demonstrated that it is possible to monitor the order and progress of the successive subroutines that are executed by the monkey by recording from area V1. The findings also illustrate how information can be transferred from one elemental operation to the next. For example, in the search-then-trace task, the initial visual search enhanced the representation of the target marker as the beginning of the target curve (at a latency of 159ms, light blue circle in Fig. 1B) so that the tracing operation could spread activity along this curve, beginning at this marker (after 229ms, light yellow in Fig. 1B). In the trace-then-search task, the trace operation enhanced the responses elicited by the target marker (at 180ms, Fig. 1C) so that its color could be registered as the target color for the subsequent search. The visual search caused an enhanced response at the circular eye movement target at a latency of 267ms (Fig. 1C).

An important finding is that the outcome of an elemental operation is made available as a focus of enhanced activity that may correspond to the working memory of that feature so that it can be used as the input for the next elemental operation [27]. These findings also illustrate the role of early visual cortex as a cognitive blackboard [16]. Reading and writing to the cognitive blackboard in early visual areas permits the exchange of information between elemental operations that are part of a visual routine. For example, in the search-then-trace task, the visual search operation tags the location of the target marker with enhanced activity in area V1, which corresponds to a write operation on the cognitive blackboard. This focus of enhanced activity marks the beginning of the tracing operation, starting at this position.

At a psychological level of description, the neuronal response modulations that occur during the recurrent processing phase correspond to shifts of attention, which are directed to image elements that are represented with enhanced neuronal activity [28]. Indeed, the sequence of mental operations is associated with a sequence of attention shifts. Visual search for a target shape or color in an array of stimuli leads to a shift of attention to the item that needs to be found [29]. Similarly, when subjects mentally trace a curve, they gradually spread attention across all of its contour elements [30].

Hence, there is substantial evidence that visual routines are implemented in the visual cortex through the propagation of enhanced activity during a recurrent processing phase. However, it remains unknown what architectures and plasticity rules permit the learning of such routines in the brain. A study by Tsotsos and Kruijne [31] considered many of the processes that are needed to implement visual routines, but they also left many of the proposed functions, like those that determine order of processing steps, for future work.

To investigate how curve-tracing can be learned, Brosch et al. [32] developed RELEARNN. RELEARNN is a biologically plausible learning rule that uses two factors to gate Hebbian plasticity. The first factor is a reward-prediction error that is central in many approaches to biologically inspired learning [33] and that is thought to be made available throughout the brain by the release of neuromodulators such as dopamine [34,35]. The second factor used by RELEARNN is a feedback signal initiated by the selected action, which propagates through a dedicated set of recurrent connections in an “accessory network” or “credit assignment network” that highlights the synapses that have had most influence on this action and that gates the plasticity. The combination of these two factors gives rise to a learning rule that is equivalent to the Almeida-Pineda algorithm that can be used to train recurrent networks [36,37], while all the information needed to control plasticity is available locally at every synapse. Importantly, RELEARNN is based on trial-and-error learning and does not require the “teacher” of error-backpropagation, because the only feedback that the network receives is whether it obtains a reward.

Interestingly, Brosch et al. [32] found that the strategy that the network learned to trace curves resembled neurophysiological results in the visual cortex of monkeys. The network learned to spread enhanced neuronal activity over the target curve. Brosch et al. [32] only tested curve-tracing and the so-called pathfinder task [38,39], which requires the detection of a string of approximately colinear line elements among other line elements with random orientations (Houtkamp and Roelfsema [40] discuss differences between curve-tracing and the pathfinder task).

It has therefore remained unclear if neural networks can learn to stitch elemental visual operations into visual routines using a biologically plausible plasticity rule that is based on trial-and-error learning. In the present study we therefore set out to examine learning of visual routines by neural networks with feedforward, feedback and horizontal connections. We asked whether the network (1) would learn multiple elemental operations, (2) whether it would learn to execute the elemental operations sequentially and in the correct order and (3) how the network ensured the transfer of information from one elemental operations to the next.

We used RELEARNN [32] to train the networks on three visual routines for which neurophysiological data in monkeys exist: a curve-tracing task, a search-then-trace and a trace-then-search task [25]. We found that the networks learned all three tasks. Interestingly, the activation of units of the networks resembled the spatiotemporal patterns of activity observed in the visual cortex of monkeys. The networks learned to propagate enhanced activity along curves and to enhance activity in retinotopic areas to transfer information between subroutines. Our results show how recurrent neural networks trained with a biologically plausible learning rule can learn to propagate enhanced activity to solve complex visual tasks.

## Results

### The tasks for the model

We trained networks on three tasks. The first task is a version of the curve tracing task (Fig. 2A). This task requires only one of the elemental operations, which is curve tracing [1]. To solve it, the model had to learn to group a set of connected pixels on a grid of 15 by 15 pixels that started with a red pixel. The model had to select the blue pixel at the end of the target curve, composed of green pixels, for an eye movement. We did not simulate the eye movement itself, but we considered the response of the network to be correct if it selected the location of the appropriate blue pixel in the output layer (Fig. 3).

**Fig. 2.**
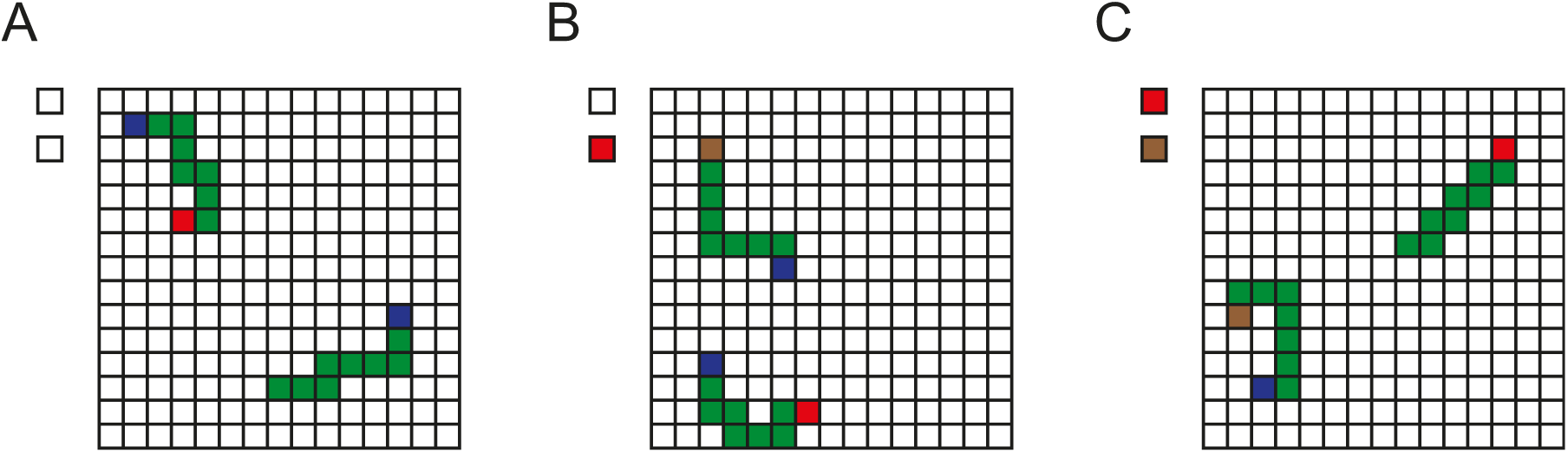
Example stimuli for the three tasks for the model. **A,** Trace task. The task is to make an eye movement to the blue pixel of the curve starting with a red pixel. **B,** Search-then-trace task. The model searches for a target marker (here red, as cued on the left of the array) and it has to make an eye movement to the blue pixel at the other end of the curve starting with this marker. **C,** Trace-then-search task. The model traces the curve starting with the blue pixel to identify a colored marker at the other end (here brown), which cued the target color that the model should select at the left of the grid. Each trial started with the full stimulus in view.

**Fig. 3.**
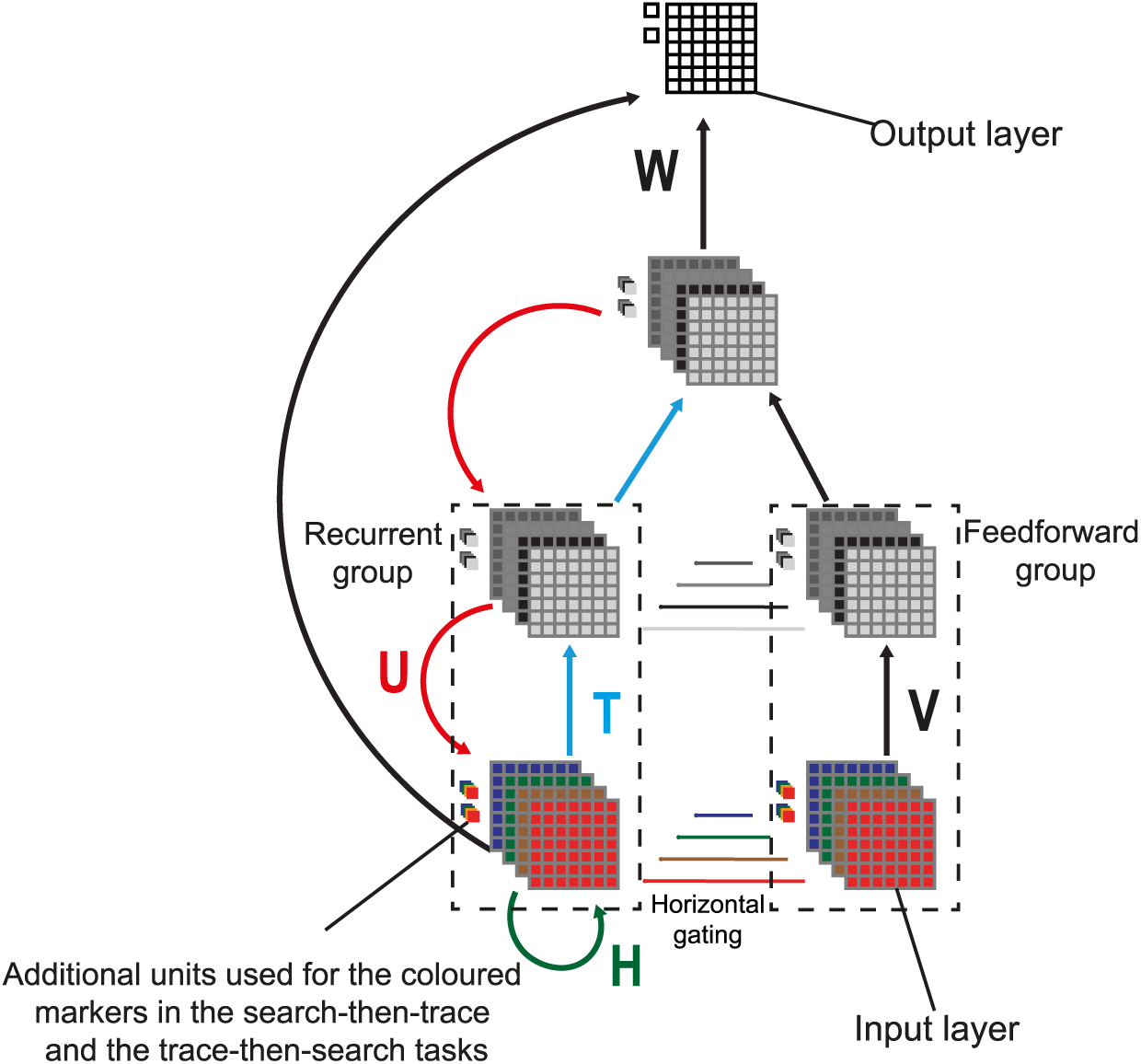
Architecture of the network. The network comprises one input layer, two hidden layers and one output layer. The input and hidden layers have four features each. In each layer, units belong to the feedforward or to the recurrent group. The activity of units in the recurrent group is gated by input of neurons in the feedforward group with the same RF so that they cannot participate in the spread of enhanced activity in case the corresponding feedforward unit is inactive.

The second task was the search-then-trace task (Fig. 2B). We cued the color that was the target of search by activating one of two additional input units, which coded for brown and red (Fig. 2). The model had to locate the pixel with the same color in the visual stimulus and trace the curve starting at this pixel to its other end to locate a blue pixel that was the target for an eye movement.

The third task was the trace-then-search task (Fig. 2C). Now the target curve started with a blue pixel. The model had to identify the color at the other end of the target curve and to select the same color at a location outside the pixel frame. Specifically, it could choose between a red and brown pixel.

In the previous experiments with the monkeys, the trial started when the animal directed its gaze to the fixation point. However, in our modeling work, we started each trial by presenting the full stimulus, irrespective of the model’s gaze position. Furthermore, we placed the colored markers that were part of the visual search subtask next to the grid of the curve tracing task, to simplify the routines. This stimulus design neither altered the structure of the tasks, nor the need to transfer information between subroutines.

The search-then-trace and the trace-then-search tasks can be solved by visual routines, composed of visual search and curve tracing as elemental operations, but in different orders. These tasks also illustrate how the output of one operation needs to be carried over as input for the next operation. For example, the output of the search operation of the search-then-trace task is the location of the marker with a specific color, which is the starting point for the subsequent tracing operation.

### Model

We will first describe the network architecture and RELEARNN, the biologically plausible reinforcement learning rule. We will then describe the network simulations and compare them to neuronal activity in the visual cortex of monkeys that were trained on the same tasks.

### Structure of the network

We trained convolutional neural networks with an input layer, two hidden layers and an output layer. The input layer (layer 0) consisted of a grid of 15×x15 input units and there were two extra input units outside the grid. There were four units at every grid location, one for each of the possible colors of the pixels (red, brown, blue and green). We did not model the more elaborate receptive field structure and its variety in low-level visual cortical areas, which might have distracted from the purpose of the model [41]. In each layer there was a group of feedforward units that only propagated information to the next layer and to other feedforward units. These feedforward units also gated the activity of units in another, recurrent group with a RF at the same position (Fig. 3, “horizontal gating”). Units in the recurrent group could propagate information to the next or previous layer and to units with different receptive fields within the same layer. As a result, units in the recurrent group could be modulated by activity outside their RF. The gating by units in the feedforward group with the same RF (Fig. 3) ensured that the response modulation could not spread to recurrent units that are not active because they represent features that are not present in the image.

The activities of neurons in the feedforward group in layer *l* at timestep *t* were determined by activities of feedforward neurons in the lower layer at the same timestep through connections *V*. If *X* represents the activity of neurons in the feedforward group, *Y* the activity of neurons in the recurrent group, *l* the layer index, then the activity of neurons was determined by:

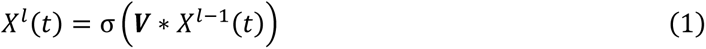

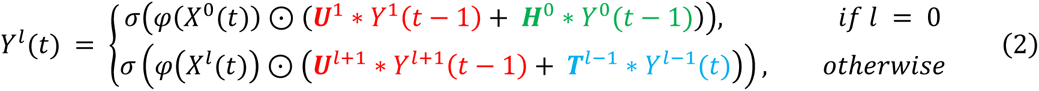

Here σ is a nonlinear activation function and φ a gating function (see below), ⊙ represents an element-wise product between the activity of feedback and feedforward units and ∗ indicates a convolution.

In the recurrent group, units in the input layer receive horizontal input from the same layer at the previous timestep through connections *H* (green terms in eq. 2), units in the higher layers receive feedforward input from the preceding layer at the same timestep through connections *T* (blue terms) and feedback from the next higher layer through connections *U* (red terms). The horizontal connections in the input layer carry color information and facilitate the learning of the search component in the search-then-trace and the trace-then-search tasks.

For connections between hidden layers, we used a 3×3 convolutional kernel, and enforced that neurons could only make connections to their Von Neumann neighborhood (4-neighborhood). As is common for convolutional neural networks, the kernels were replicated using weight sharing, a property not present in biological neural networks [42]. However, the networks could also learn the tasks without weight sharing. Specifically, without weight sharing the networks learnt the curve-tracing task after an average of 140,000 trials as opposed to 19,000 trials when we used weight sharing. We used weight sharing to reduce computation time, but we note that our results are likely to generalize to biologically plausible learning schemes without weight sharing.

The last hidden layer was fully connected to the output layer and the recurrent input layer with weights linking units with the same RF initialized with positive values and between units with different RFs initialized with negative values to bias the winner-takes-all mechanism in the last layer, which stabilized learning.

The activity of units in the retinotopically organized output layer was determined as:

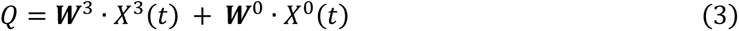

where *T* is the timestep at which the network reached a stable state (Fig. 3). Units in the output layer learned to represent the also called Q-value [33], which is the expected reward when the networks selected this position for an eye movement.

During training, the model chose the eye movement with the highest Q-value with probability 1 − *∈*. With probability *∈* the network explored other actions and sampled a random action *a* from the Boltzmann distribution *P*_*B*_:

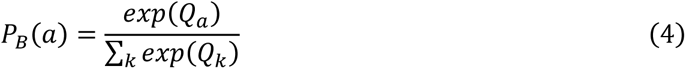

*∈* was set to 0.05 across tasks and networks. As mentioned in the above, we only simulated the selection of the eye movement in a map of space. We neither simulated the eye movement itself nor the shift of the visual image that is caused by eye movements.

We trained the networks with RELEARNN [32], which is a biologically plausible implementation of Q-learning that uses three phases. Upon presentation of the stimulus, the activity of the input neurons remains constant and activity propagates through the recurrent connections of the network. We selected the action once the activity of the units settled, but we did not wait longer than 50 timesteps. When an action was selected, an “attentional” signal originating from the winning action was propagated through an accessory network to determine the influence of each neuron on the selected action. The accessory network is important for credit assignment, with one unit for every unit of the regular network. It permits local computation of weight updates (see “RELEARNN”). The network then gets a reward *r* from the environment and computes a reward prediction error δ that is determined by:

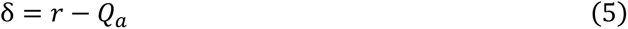

Where *Qa* is the Q-value of the selected action *a*. The reward prediction error δ is broadcasted to the whole network by a neuromodulatory signal, such as dopamine [34] and is thus available at all synapses. δ and the “attentional” feedback signal are combined to update the weights of the network, as will described below in the RELEARNN section.

### Activation and gating functions

A popular activation function to train recurrent neural networks is the ReLU function because it alleviates the problem of the vanishing gradient [43]. However, ReLU functions offer no guarantees that the network will reach a stable state [44], which is a necessary condition for learning rules such as RELEARNN. On the other hand, squashing nonlinearities like sigmoid or tanh functions guarantee that the network reaches a stable state [45], but may suffer from the vanishing gradient problem, which might hurt performance [46]. To guarantee that our network reaches a stable state while mitigating the vanishing gradient problem, we used a ReLU-like function with a slope that decreases after a threshold:

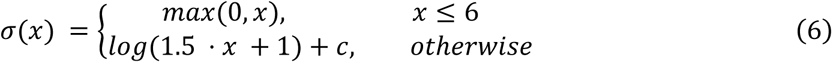

Where c is set to 6 − *log*(10) to ensure that *σ* is continuous. We will describe below how the networks learn to label relevant pixels with a response enhancement. We observed that this process was hampered if the activation function was bounded from above when curves were long. In that situation, units coding for all pixels reached the bound and the networks could no longer differentiate between the target and the distractor curve. The activation function of Eq. 6 ensured that the network differentiated between long target and distractor curves.

The horizontal gating of feedback units by feedforward units (see eq. 2) was also governed by a non-linear function. The activity of neurons of the feedforward units was passed through a continuous approximation of a step-function:

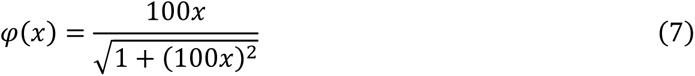

Here *φ*(*x*) is non-negative because *x* is non-negative (see Eq. 2 for the role of *φ*(*x*)).

## RELEARNN

A popular algorithm to train recurrent neural networks is backpropagation trough time (BPTT) [47,48]. The algorithm needs to memorize the consecutive states of every unit because this information is needed to update the weights. It is non-biological because it is neither local in time nor in space [49]. We therefore used RELEARNN [32], a learning rule that is local in time and in space and equivalent to backpropagation in a reinforcement learning paradigm [36,37]. RELEARNN uses a separate accessory “attentional” network with one accessory unit for every regular network unit (Fig. 4). On each trial, the output layer chooses one of the possible actions. The accessory units represent the influence of the corresponding regular units on this chosen action. Once an action has been chosen, the winning unit in the output layer activates accessory units and thereby initiates the propagation of activity in the accessory network. The strength of a connection from unit A to unit B in the accessory network is proportional to the strength of the connection from unit B to unit A in the regular network so that the activity in the accessory network (dashed red and green lines in Fig. 4) equals the influence of regular units on the chosen action, once the accessory network converged to a stable state (Fig. 4, see Brosch et al. [32] for details). Accessory units corresponding to regular units with a strong influence on the Q-value of the chosen action are very active and those that correspond to regular units that did not influence this Q-value are silent. The network of accessory units can thereby assign credit (or blame) to the regular units that are responsible for the outcome of the action. A change in the synapses between the regular units with highly active accessory units has a pronounced influence on the Q-value of the chosen action.

**Fig. 4.**
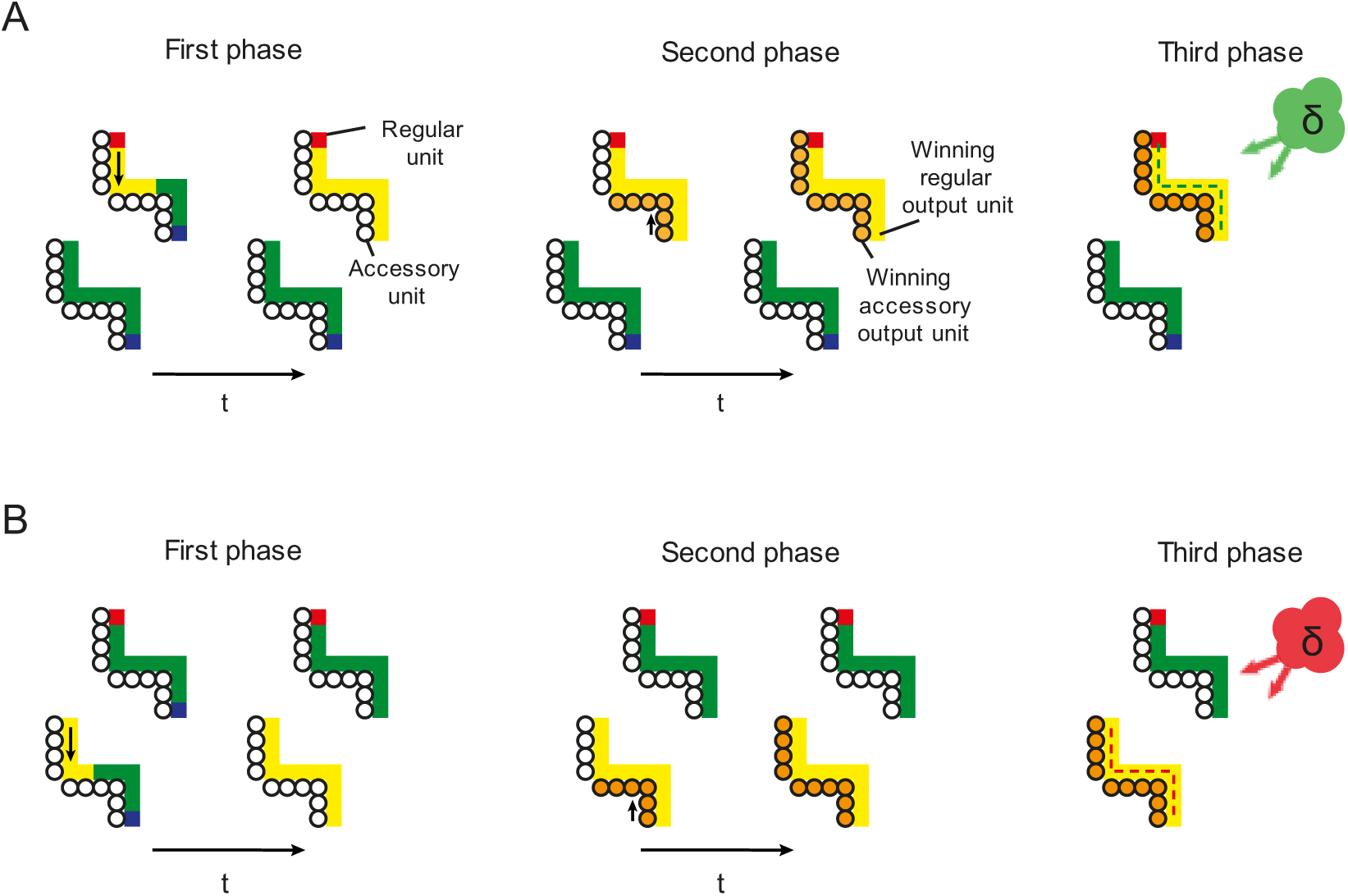
RELEARNN for an example stimulus. **A. Correct choice by the network.** In the first phase, activity propagates in the regular network that has both feedforward and recurrent connections (squares) until this network reaches a stable state. Here enhanced activity (yellow) spreads over the target curve. In the second phase, a winning unit is selected and activates the corresponding unit of the accessory network (small circles). From there, activity propagates in the accessory network (small orange circles) to tag the connections that influence the Q-value of the chosen action. After a few timesteps, activity in the accessory unit 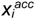 becomes proportional to the influence of the activity of the corresponding regular unit 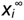 on the chosen output unit *Qa*. In the third phase, a reward is given if the action was correct, or not in case of an error, and a neuromodulator (green cloud, δ) broadcasts the reward prediction error to the network. Weights are changed according to a four-factor Hebbian learning rule (green connections between the units are increased). **B. Incorrect choice by the network**. In this case, the enhanced activity spreads over the wrong curve and reward prediction error is negative because of the wrong choice (red cloud). Hence, the weights between units that represent the distractor curve are decreased (red connections).

We distinguish between three phases on a trial (Fig. 4). In the first phase, the stimulus is presented as a pattern of activity across the input neurons. Activity propagates trough the feedforward, horizontal and feedback connections among regular feedforward and recurrent units until the network converges to a stable state. At the end of this phase a presynaptic neuron *i* and a postsynaptic neuron *j* will have activity 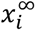 and 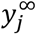. At the start of the second phase, an action is selected, which is represented by one of the regular units in the output layer. The corresponding unit in the motor layer of the accessory network is activated and activity is propagated within the accessory network until a stable state is reached (in our experience this phase can be terminated after a fixed number of timesteps [34]). At the end of the second phase, neurons *i* and *j* in the accessory network have activity 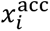 and 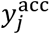, which is proportional to the influence of the corresponding regular units on the chosen action value (see Brosch et al. [32] for a proof):

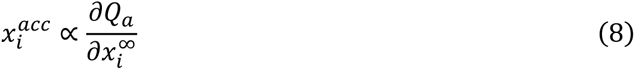

In the third phase, the network receives a reward if the correct action was selected, and no reward in case of an erroneous response. The reward prediction error is broadcasted through the network using a neuromodulator and weights are modified according to a four-factor Hebbian learning rule:

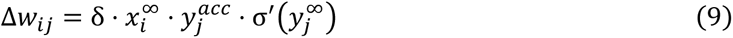

where Δ*w*_*ij*_ represents the change in weight *w*_*ij*_, δ is the reward prediction error defined in Eq. (5), 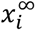 and 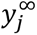 are the activities of neurons *x*_*i*_ and *y*_*j*_ in the main network when a stable state is reached and 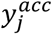 is the activity of neuron *y*_*j*_ in the accessory network once this network has also reached a stable state. Note that the weight update of RELEARNN is local in space, as weights are modified using only pre- and postsynaptic information, and local in time, because it is not necessary to memorize the activities of the units at every time step.

### Curve-tracing task

We trained the networks with a curriculum to perform the curve-tracing task, a strategy that is also used to train monkeys. We first presented a stimulus with one blue and one red adjacent pixel, which were placed randomly on the grid, to train the network to select blue pixels as target for an eye movement. If the network achieved an accuracy of 85% during a test phase with 500 trials, during which we fixed the weights and switched exploration off, the network was trained with a target and a distractor curve with a length of three pixels. When the network achieved an accuracy of 85%, we added one pixel to the curves until the length of the curves was 9 pixels. Using this curriculum, we trained 24 networks and judged that they had learned the task if they achieved an accuracy larger than 85% for curves that were 9 pixels long. All networks learned the task, within an average of 19,000 trials. By comparison, monkeys trained on curve-tracing tasks take weeks to months with thousands of trials per day to learn the task. The total number of trials they undergo during training hence adds up to the tens of thousands.

We next aimed to gain insight into the strategy of the networks. The curves were drawn randomly on each trial (see Fig. 5A for an example stimulus). The number of possible curves grows exponentially with their length, and we therefore conjectured that it might be impossible to memorize all possible input patterns [50]. Instead, we hypothesized that the networks learned a general rule, which could be based on applying an elemental curve-tracing operation. To test this hypothesis, we exposed the networks to curves that were longer than those shown during training. When a network achieved an accuracy larger than 85% for curves of length N, we switched off exploration and tested curves with lengths ranging from N+1 to N+4. We found that the networks’ accuracy was above chance levels for curves that were longer than the curves in the training set (Fig. 5B). For example, networks trained on curves up to a length of 9 pixels generalized to curves with a length of 13 pixels (red in Fig. 5B) (p<10^−6^, Wilcoxon signed-rank test). This generalization emerged during training, indicating that networks learned to trace connected pixels as a generalized elemental operation [50].

**Fig. 5.**
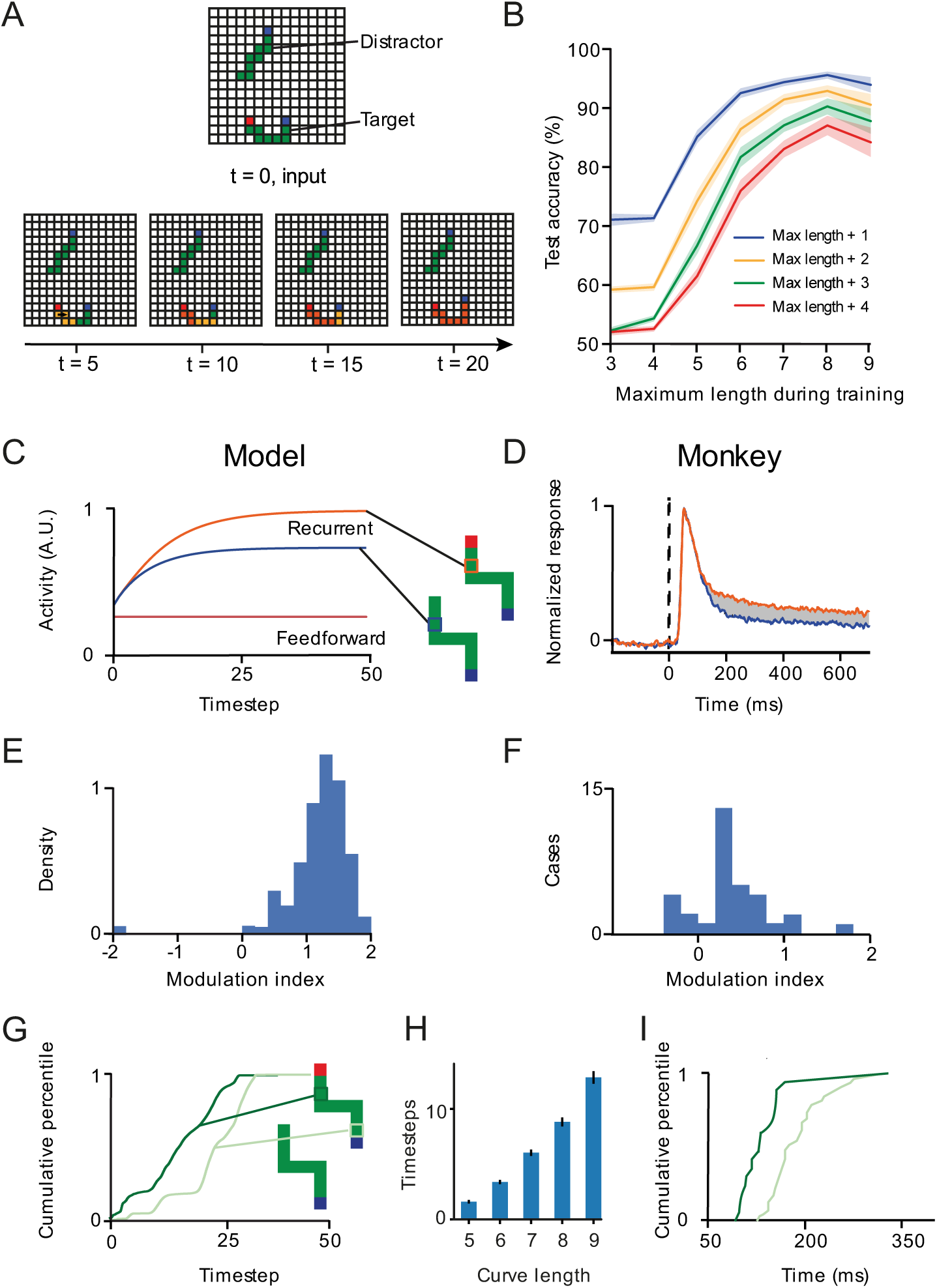
Propagation of enhanced activity across the representation of the target curve during curve-tracing. **A**. Upper, example stimulus presented to one of the networks. The target curve starts with a red pixel. Lower, activity of recurrent units in the input layer across time. The orange color denotes an increase in activity. Note the spread of enhanced activity over the representation of the target curve, starting at the red pixel. **B**. Testing accuracy for curves of length up to N+4 pixels where N is the maximum length used during training. At the beginning of training, the model does not generalize to longer curves. At the end of training, a model trained with curves up to 9 pixels long generalized to curves with up to 13 pixels (p<10^−6^, Wilcoxon signed-rank test). **C**. Activity of an example unit in the recurrent group elicited by the target (orange) or distractor curve (blue), and activity of the corresponding unit in the feedforward group (brown). The activity elicited by the target curve is enhanced compared to that elicited by the distractor curve**. D**. Average activity of neurons in area V1 of the visual cortex of monkeys during a curve tracing task, when their RF fell on the target curve (orange) or on the distractor curve (blue). Adapted from (Roelfsema et al., 2003). **E.** Distribution of the modulation index across recurrent units of the neural networks. A positive value indicates an enhanced response to the target curve. **F**. Distribution of modulation index in area V1 of the visual cortex of monkeys (from Roelfsema et al., 1998) **G**. Distribution of the modulation latency across units of the network. The onset of modulation is delayed for units representing pixels that are farther (7 pixels away), compared to pixels that are closer (2 pixels away) to the beginning of the curve (p<10^−15^, Mann-Whitney U test). **H**. The minimum number of timesteps needed to reach 85% accuracy increased for longer curves, indicating the need for recurrent processing. Error bars, 95%-confidence intervals. **I.** Distribution of the modulation latency across recording sites in monkeys performing the curve-tracing task, adapted from Pooresmaeili & Roelfsema, 2014. Dark green represents RF that were close to the fixation point, and light green represents RF that were farther from the fixation point.

We next examined the activity of units of trained networks and compared it to the activity in the visual cortex of monkeys that had been trained on a similar curve tracing task [17]. We first compared the activity of networks units whose receptive field fell on the target curve to the activity of units whose receptive field fell on the distractor curve. By design, the activity of the units of the feedforward group is not modulated during curve-tracing, whereas units of the recurrent group could learn to propagate enhanced activity (Fig. 5A,C). Indeed, the activity of units whose RF fall on the target curve is enhanced compared to units whose RF fall on the distractor curve.

Fig. 5A illustrates the flow of activity in the recurrent input layer of one of the networks for an example stimulus. At t=0, the stimulus is presented, and the feedforward flow of activity is complete. Feedforward units with a pixel of the appropriate color in their receptive field are active but the recurrent units do not yet discriminate between the target and distractor curve. After a few timesteps, enhanced activity starts to spread from the red cue at the beginning of the target curve. However, the end of the curve is not yet labeled, and the network doesn’t have enough information to make the appropriate eye movement. After 40 timesteps, however, the enhanced activity has spread over the entire representation of the target curve to reach the blue eye movement target, which is selected by the network.

The gradual spread of enhanced activity across the target curve resembles the spread of enhanced neuronal activity in the visual cortex of monkeys solving a similar curve-tracing task. Fig. 5D shows the average activity of V1 neurons in the visual cortex of monkeys elicited by a target curve (orange) and a distractor curve (blue). Initially, the neurons were activated by the appearance of a contour element in their RF, and this feedforward response was the same for the target and distractor curves. After a delay of 130ms, recurrent processing caused the V1 representation of the target curve to be enhanced over the representation of the distractor curve [26].

Once the training of 24 networks had completed successfully, we used a modulation index [11] to quantify the magnitude of the response enhancement across 345,000 stimuli. The modulation index represents is the difference between the response evoked by the target and distractor curve divided by the average response:

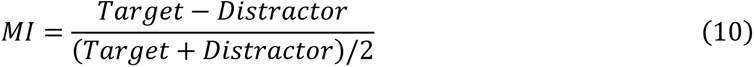

A positive modulation index signifies an enhancement of activity evoked by the target curve relative to that evoked by the distractor curve. We observed that the recurrent units representing target pixels had a higher activity than those representing distractor pixels. The mean modulation index was 1.18, corresponding to an activity enhancement of 292% (Fig. 5E). Neurons in area V1 of monkeys that responded to the target curve also enhanced their activity, although the average activity increase was weaker (Fig 5F, see also Roelfsema et al., 1998). We note, however, that the V1 modulation index distribution in the monkeys included neurons that were not modulated by curve-tracing, resembling feedforward units of the network.

In the visual cortex of monkeys, the enhanced activity spreads iteratively, starting at the red cue at the beginning of the curve until it reaches the end (Fig 5H). Accordingly, units in the model representing pixels close to the beginning of the target curve enhanced their activity before units representing pixels that were farther away. To quantify the time-point of modulation, we used the moment when the difference between the activity elicited by the target and distractor curve became larger than 70% of its maximum (Fig. 5G). An analysis of modulation latencies across units confirmed that the onset of modulation was significantly later for pixels farther from the fixation point (7 pixels away), compared to pixels closer to the fixation point (2 pixels away) (p<10^−15^, Mann-Whitney U test). The reliance on the spread of activity was confirmed by examining the minimal number of time steps that the networks needed to solve the task with curves ranging from 5 to 9 pixels (Fig. 5H). The number of timesteps needed to reach 85% accuracy increased with the length of the curves. For example, the minimal number of time steps was 2 for curves with a length of 5 pixels and increased to 13 for curves with a length of 9 pixels. These results confirm that recurrence is needed to solve the task. Interestingly, the networks could also trace longer curves including spirals with up to 25 pixels (Fig. 6A) and stimuli with many distractors (Fig. 6B,C), further confirming the generality of the solution.

**Fig. 6.**
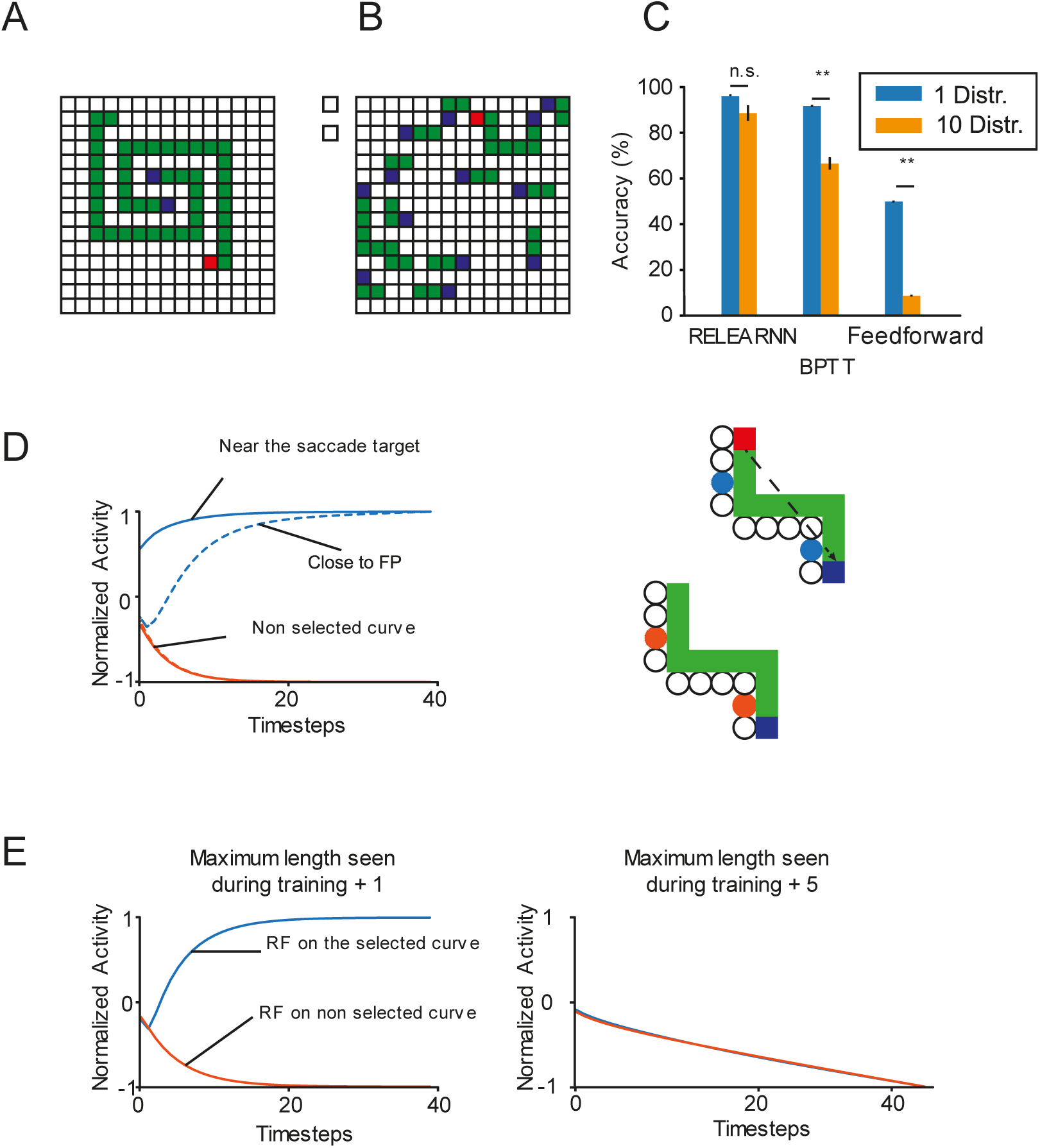
RELEARNN mechanisms. **A,B.** More challenging curve tracing stimuli with long spirals (A) or with many distractors (B). **C**. Accuracy of networks trained on the curve-tracing task with one distractor, when tested on the curve-tracing task with 10 distractors. The networks trained with RELEARNN could solve the task as well, irrespective of the number of distractors (p=0.17, Mann-Whitney test). Networks trained with BPTT did not generalize as well (p<10^−5^, Mann-Whitney test) and feedforward networks could not be trained on the curve-tracing task, i.e. they were at chance level. **D.** Activity of units in the accessory network whose RFs fall on the selected curve (blue traces) or the non-selected one (orange traces), at different distances from the blue pixel that is target of the eye movement (continuous and dotted traces show the activity of accessory units representing pixels nearer to and farther from the saccade target, respectively). Hence, the credit assignment signal propagates in the opposite direction than to the enhanced activity, starting from the selected eye-movement target. This credit assignment signal is absent from the representation of the distractor curve. **E.** Activity of units at the beginning of the selected and non-selected curves in the accessory network, for curves that were one (left panel) or five pixels longer (right) than the curves used during training. If the length of the curve was similar to that in the curriculum, the credit assignment signal propagated to the beginning of the selected curve (red fixation point on correct trials) and training is effective. However, if the curves are much longer, the credit assignment signal does not spread to all other pixels of the selected curve and training fails.

In monkeys, the spread of activity among neurons in the visual cortex is accompanied by an enhancement of the activity of neurons coding for the target curve in the frontal eye fields, which is an area involved in the planning and generation of eye movements [51,52]. Interestingly, the spread of enhanced activity was also evident in the model’s output layer, which represents the selected eye movement (Fig. S1). Hence, the model had learned a strategy of spreading enhanced neuronal activity along the entire target curve in multiple layers. In the model, spreading is iterative and relies on horizontal and feedback connections, which makes the process generalizable across stimuli and curve lengths, in spite of the limited network depth. The results are also in line with the results by Jolicoeur et al. [5] showing that the reaction times of human observer in a curve-tracing task increase linearly with the length of the curve that needs to be traced.

We examined whether the task could be solved by a feedforward network with the same architecture, with 3×3 kernels. However, we found that these feedforward networks did not solve the task for curves longer than 3 pixels. In previous work [50], we observed that feedforward networks need to reserve a hidden unit for every possible shape of the target curve, and this strategy presumably failed, given the large variety of curves.

How did the network learn to trace, i.e. to group pixels into curves, when the only feedback after each trial is the presence or absence of a reward? To understand the learning process enabled by RELEARNN, we examined the activity of the accessory network units that influence learning. Figure 6D illustrates how the activity propagates among the accessory units that represent the selected curve, starting from the selected eye movement and then across the representation of other pixels of this curve, i.e. in the opposite direction of the activity propagation. The strength of this accessory signal represents the influence of the units of the regular network on the selected action. If the network happens to select the target curve, the connections between regular units are strengthened in proportion to the reward-prediction error δ. If the network erroneously selects the distractor curve, the credit assignment signal ensures that the recurrent connections among units representing this curve are weakened because δ is negative. The unique and distinguishing feature of the target curve is that it starts with a red pixel, and the credit assignment signal of the accessory network ensures that the representation of this red pixel becomes the source of activity propagation in well-trained networks during learning.

The analysis of the spread of activity within the accessory network also provided insight in the role of the curriculum. If the network has been trained with curves up to a certain length N, and is tested with curves that have N+1 pixels, the credit assignment signal reaches the fixation point if the model selects the appropriate eye movement and the relevant connections increase in strength (Fig. 6E, left). However, when we presented curves with N+5 pixels, the signal at the beginning of the curve did not discriminate between the target and distractor curves and learning did not occur (Fig. 6E, right). Hence, training was more efficient with a curriculum in which we gradually increased the length of the curves.

RELEARNN provides a biologically plausible approximation to BPTT [32] and we trained 24 networks with BPTT to perform the original curve-tracing task and tested generalization to the version with many distractors. The architecture of the networks trained with BPTT was the same as those trained with RELEARNN and their accuracy was reduced for the stimuli with many distractors (Fig. 6B). This drop in accuracy was caused by the proximity between the red pixels of the target curve and blue pixels of nearby distractors, which were erroneously selected for an eye movement. This drop in accuracy did not occur for networks trained with RELEARNN, indicating that RELEARNN caused better generalization to stimuli not shown during training.

Let us briefly summarize how the networks learn to solve the curve-tracing task. Upon presentation of the stimulus, the network starts with a feedforward processing phase which does not discriminate between the target and the distractor curve. This early activity pattern has been called ‘base representation’ in previous work [15]. During the later, recurrent processing phase, units learn to propagate enhanced activity along the representation of the target curve, starting at the fixation point with its unique color. This enhancement of activity cannot spill over from the target curve to nearby curves because feedforward neurons that respond to image locations without pixels are silent. The activation of these feedforward neurons is necessary before the recurrent units activate. In other words, the gating process of the base representation ensures that enhanced activity can only spread along the representations of connected pixels.

### Search-then-trace task

Having established that the networks learn to trace curves, we explored more complex multistep visual routines that require the succession of multiple elemental operations, and the transfer of information between them. The search-then-trace task required a visual search for a colored marker followed by curve-tracing. In the task for the model, the color of relevant markers was brown or red and the color of the target marker was cued by activating one of two input units (color cue in Fig. 7A). The network had to trace the target curve starting at this marker and to select an eye movement to the blue pixel at the other end of this curve.

**Fig. 7.**
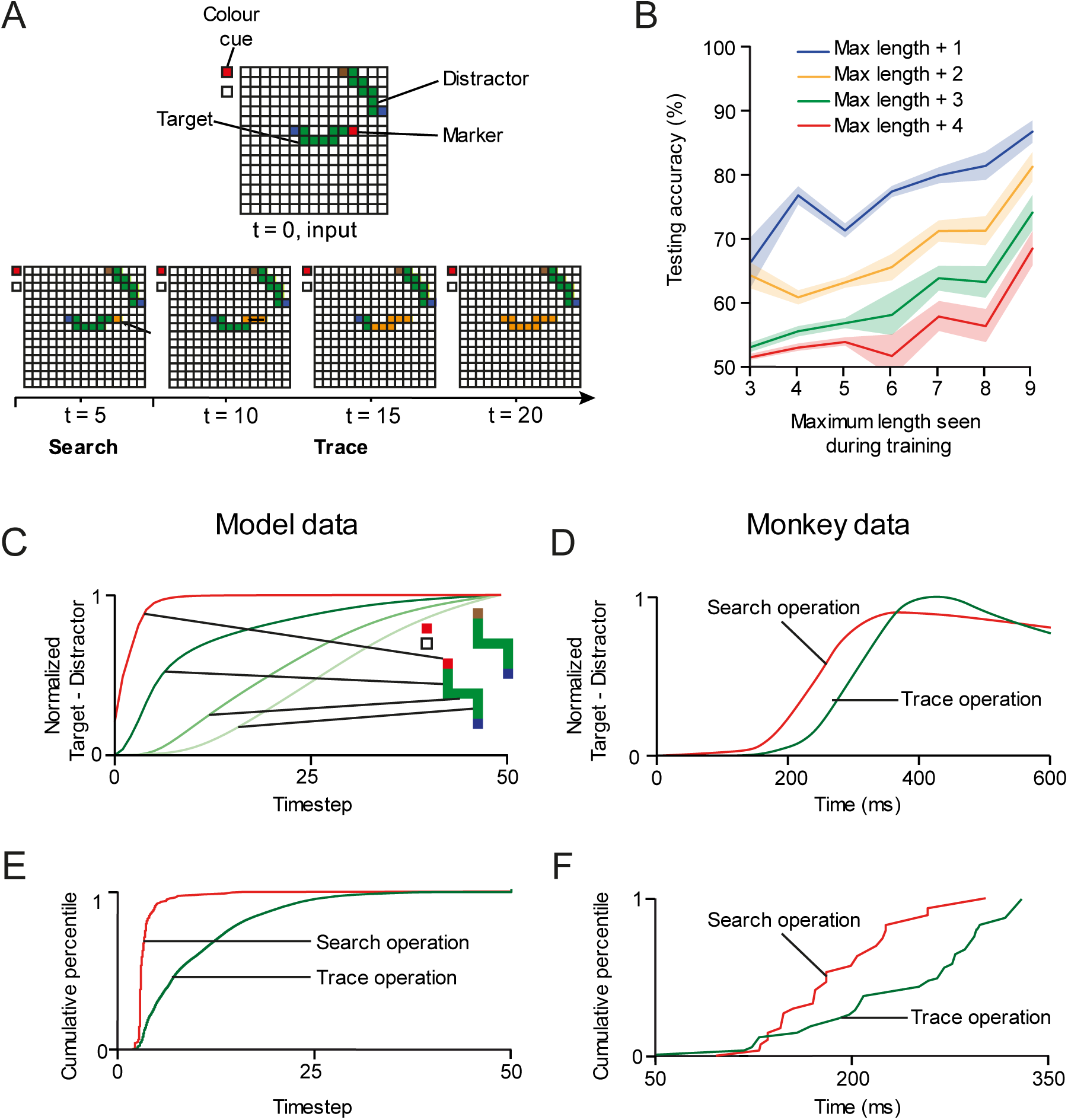
Search-then-trace task. **A.** Example stimulus shown to one of the networks. Upper panel, visual stimulus. Lower panel, orange shading shows the propagation of enhanced activity among recurrent units of the input layer, starting at the representation of the red marker, which is highlighted as the result of the search operation. From here, the enhanced activity spread along the curve (trace operation). **B.** We tested how well the models generalized to curves that were longer than those presented during training. Generalization was better for networks that had been trained on longer curves (x-axis). E.g. networks trained on curves up to a length of 9 pixels generalized to curves with 13 pixels (p<10^−6^, Wilcoxon signed-rank test). **C**. Normalized response enhancement for the target marker and target curve. Each curve is normalized by its maximum over time. First the activity of the unit with a RF at the location of the target marker was enhanced (search operation, red curve). Thereafter, enhanced activity propagated across the target curve connected to it (trace operation, green curves). **D**. In the visual cortex of monkeys, the representation of the target marker is enhanced (red) before the enhanced activity spreads over the V1 representation of the target curve (green; adapted from Moro et al. 2010). **E**. Distribution of the latency of the response enhancement across 260,000 stimuli and 19 networks. The latency of the modulation related to the search operation was shorter than that related to curve-tracing (p<10^−15^, Mann-Whitney U test). **F**. Distribution of the latency of response enhancements across V1 neurons in monkeys solving the search-then-trace task (adapted from Moro et al., 2010).

We trained the network using a curriculum that was similar to the one used for the trace task. First, we presented a stimulus with a blue pixel connected to a red pixel randomly placed on the grid. If the network achieved 85% accuracy during a 500 trials test phase with weights fixed and no exploration, we presented a target and a distractor curve with a length of three pixels, where one of the two curves started with a brown pixel and the other one with a red pixel. If the network achieved 85% accuracy during the test phase, we added one pixel to the curves and repeated this until the curves were 9 pixels long.

We trained 24 networks and 79% of them learned the task within an average of 40,000 trials, which is less than the number of trials that monkeys performed during weeks of training before they could do the task. Just as in the curve-tracing task, the networks learned a solution in which they generalized across different shapes of the curves and for both marker colors. Trained networks generalized to curves that were 4 pixels longer than the ones presented during training (Fig. 7B, red curve, p<10^−4^, Wilcoxon signed-rank test).

How does the network learn to transfer information between elemental operations such that the output of an operation can be used at the input of the next one? In the visual cortex of monkeys, this information transfer appears to take place by the persistence of enhanced neuronal activity in early visual cortex between the elemental operations. In the search-then-trace task, the visual search for the marker with the cued color enhances the activity of neurons with a RF at this marker (Fig. 1A), as if it is ‘written’ to a cognitive blackboard represented in early visual cortex [4,16,53]. he subsequent tracing operation can ‘read’ this position and use it as the start for the curve-tracing operation.

Interestingly, the model discovered a similar method to transfer information between the successive search and trace operations. In the model, recurrent processing started with an enhancement of the representation of the target marker in the recurrent input layer (Fig. 7A). After training, the horizontal connections in the recurrent input layer between the color cue units and the retinotopic units of a same color are strengthened compared to those between the other colors, and the enhanced activity propagated through those connections to the retinotopic unit with the same color as the cue. This strategy solved the search operation.

As in the trace operation, the gating by the feedforward units of color cue units and retinotopic units ensured that the enhanced activity due to the search operation is confined to units that represent features that are present in the stimulus.

After the search operation, the enhanced activity propagated across the representation of the target curve (Fig. 7A, green traces in Fig. 7C). As in the curve-tracing task, the time course of activity propagation in the output layer was similar to that in the visual layers of the model (Fig. S1). We next examined the time course of enhanced activity of the units across 260,000 stimuli and the 19 networks that learned the task. This analysis revealed that the early enhancement of the activity of units with a RF on the target marker (red in Fig. 7E), followed by the propagation of enhanced activity across the target curve (green in Fig. 7E) was a general feature. Fig. 7F shows similar results in area V1 of the visual cortex of monkeys performing a search-then-trace task [25]. Also in the monkey visual cortex, the marker with the relevant curve is first labeled with enhanced activity, before activity spread across the V1 representation of the target curve.

These results, taken together, indicate how RELEARNN, a biologically plausible reinforcement learning rule, can train networks to execute a visual routine composed of multiple elemental operations and to use persistent enhanced activity to transfer information from one element operation to the next.

### Trace-then-search task

We next used the trace-then-search task to examine if the network could learn a different visual routine composed of the same elemental operations that need to be applied in the opposite order. First, the agent had to trace the curve connected to the fixation point, which was now blue. This target curve ended with a marker that was either red or brown which was the color that the model had to search because it had to select the pixel with this color on the side of the grid (Fig. 1C).

We adapted the curriculum by first training the networks on simple curve-tracing tasks in which the target curve started with a blue pixel, and the two curves ending with a red and a brown marker. When the networks had learned to trace curves with a length of 9 pixels, we added the two colored pixels outside the grid and required the model to select the pixel with the same color as the target marker for an eye movement.

Of the 24 networks trained on this task 66% learned it, after an average of 37,000 trials. As in the previous tasks, the networks learned a general solution, which did not depend on the position or color of the marker or on the shape of the curve. Trained networks also performed the task for curves that were 4 pixels longer than the curves of the curriculum (Fig. 8B, p=1.4·10^−4^ for curves with a length of 13 pixels, Wilcoxon signed-rank test).

**Fig. 8.**
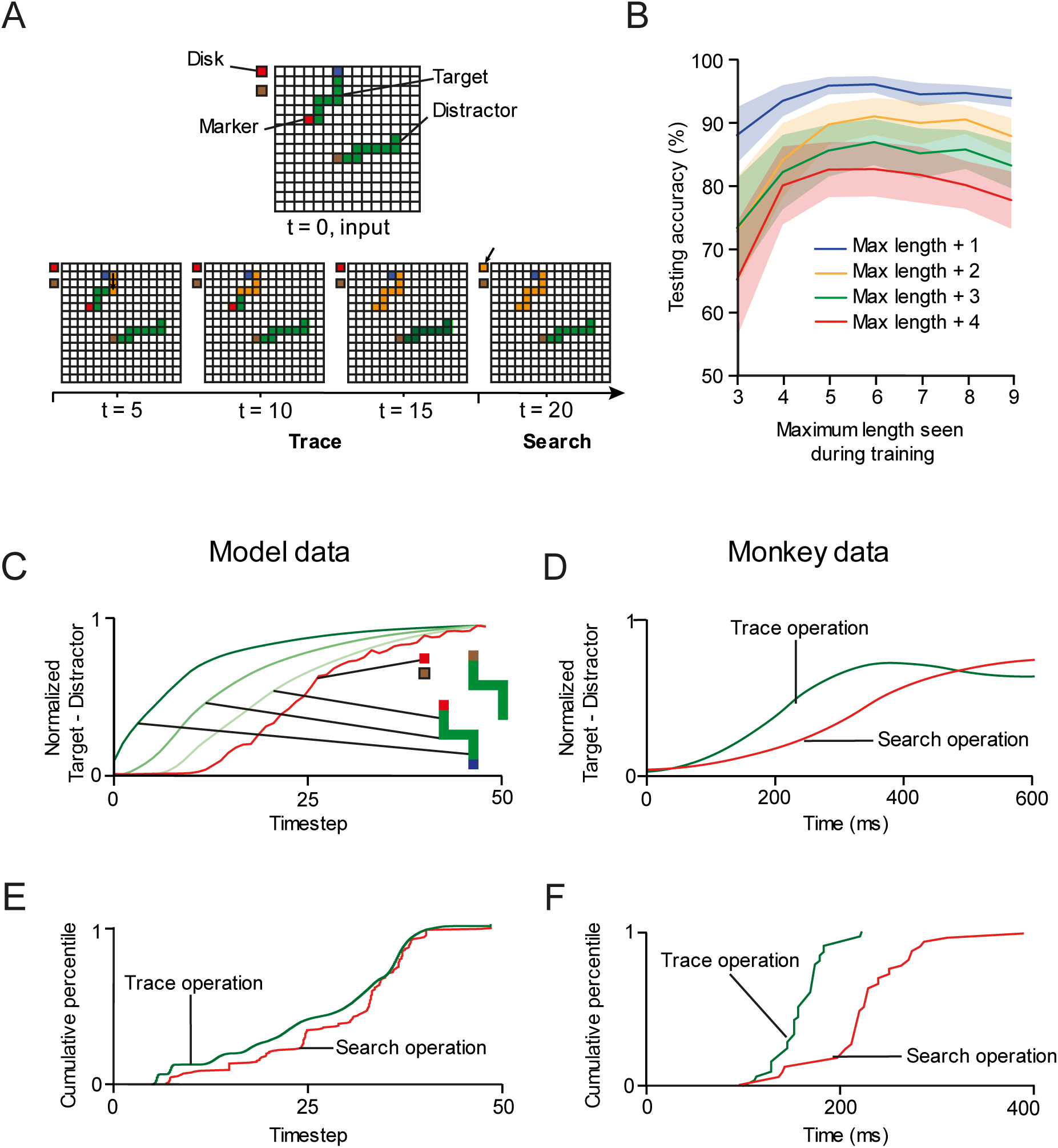
Model performance in the trace-then-search task. **A.** Example stimulus shown to one of the networks. Upper, an example stimulus. Lower, the spread of enhanced activity is shown in orange. It first spreads over the curve starting at the blue cue and reaches the target marker at the other end, cuing the color that needed to be selected during the search operation. **B.** Testing accuracy for curves of length up to N+4 pixels where N is the maximum length in the curriculum. The generalization performance improved when the network learned to trace longer curves (p=1.5·10^−4^ for curves of 13 pixels, Wilcoxon signed-rank test). **C**. Normalized response enhancement for target pixels, averaged across units. Each curve is normalized by its maximum over time. First the curve connected to the fixation point is labeled with enhanced activity (trace operation, green curves) and then the units that represent the correct eye movement target, i.e. with the same color as the target marker, enhanced their activity (search operation, red trace). **D**. In the visual cortex of monkeys, the response enhancement also first labels the segments of the target curve (green trace), before it labels the position of the eye movement target (red trace; adapted from Moro et al., 2010). **E**. Distribution of the modulation latency across model units (230,000 stimuli and 16 networks). The response modulation of trace operation precedes that of the search operation (p=1.5·10^−5^, Mann-Whitney U test). **F**. Distribution of the modulation latency across recording sites in monkeys solving the search-then-trace task (adapted from Moro et al., 2010).

Trained networks first spread activity along the target curve until its end was reached (Fig. 8A,C) and the target color could be identified. The enhanced activity then spread to colored eye movement target, as the outcome of the search operation. Hence the networks learned to first perform a trace operation, followed by a search operation. As in the previous tasks, the time course of modulation in the output layer followed a similar pattern as in the input layer (Fig. S1). The progression of activity was also evident when we examined the activity across 230,000 stimuli in the 16 networks that were trained successfully. The green trace in Figure 8E illustrates the time-point in which the activity of the representation of the last, colored pixel of the curve was enhanced, which corresponds to the output of the curve-trace operation (green curve). Thereafter, the representation of the colored eye-movement target was enhanced, corresponding to the realization of the search operation (red curve in Fig. 8E). The results are qualitatively like those in the visual cortex of monkeys trained on the trace-then-search task, where the enhanced neuronal activity first spread along the target curve (Fig. 8F, green trace), before the V1-representation of the colored disk that was the target of the eye movement was enhanced (Fig. 8F, red curve).

These results indicate that RELEARNN can train networks to carry out elemental operations in different orders depending on the task at hand, which is remarkable for a biologically inspired learning rule in which the only external feedback that the networks receive about the outcome of the decision is a reward upon a correct choice.

## Discussion

We studied how visual reasoning tasks can be learned by trial and error, using a biologically plausible learning rule. We took inspiration from previous work on visual routines, which are built from elemental operations [1,5,31]. Studies of the visual cortex revealed that elemental operations, like curve-tracing and visual search, are associated with the enhancement of neuronal activity elicited by target elements over distractor elements. Specifically, curve-tracing is associated with the spread of enhanced activity across a target curve and visual search results in an activity enhancement of the representation of the target of search (see also ref. [53,54]). We found that the neural networks trained to trace curves or to carry out visual routines relying on search and curve-tracing developed a similar strategy. They learned to successively label task-relevant image elements with enhanced neuronal activity. We also studied how these networks learned to transfer information between one elemental operation and the next. We observed that the strategy of labeling image elements with enhanced activity provides a mechanism for information transfer, because the focus of enhanced activity that was the result of one operation could be used to start the next elemental operation. Note that this mechanism is equivalent to the binding of parameters to subroutines in a computer program. The networks thereby provide new insights into how the processes that are necessary for a neuronal implementation of visual routines (see ref. [31] for an inventory) can be learned by trial and error.

An important difference between the artificial neural networks trained by us and the visual system of humans and monkeys is in their prior visual experience. The humans and monkeys tested in previous work in curve tracing and visual routines knew how to segment and attend to objects, given their previous experience. Humans struggle to learn new visual tasks when such sources of prior knowledge are removed from the visual stimuli [55]. Furthermore, monkeys can only be trained to trace curves using rewards as the incentive, which takes weeks to months with thousands of trials per day. Hence, the total number of trials while training a monkey adds up to tens of thousands, which is comparable to the number of trials that were necessary for models trained with RELEARNN. The models of the present study had to learn the task structure, how to trace a curve and to search for an item of a particular color without any prior knowledge. To train these models on visual routines, we used curriculums in which the complexity of the task gradually increased, resembling the curriculums used to train monkeys on visual routines [25,26].

The model architecture included a feedforward group of units that is only sensitive to input from lower layers, and a recurrent group that also receives feedback connections from higher layers and horizontal connections from units in the same layer. Such a division of labor between feedforward and recurrent units is advantageous, because it separates attentional effects from the representation of low-level features like stimulus contrast [56], and has also been used in previous studies. Indeed, several models [54,57–59] studied neural networks with a feedforward pathway for feature detection and a feedback pathway for attentional selection. Similar observations have been made in the visual cortex, where some neurons are influenced by curve-tracing and other neurons are not [56]. The neurons that are and are not sensitive to recurrent processing are clustered in different layers of cortex, as has been observed during curve-tracing and figure-ground segregation tasks [27,60]. Neurons that are most sensitive to the feedforward input are enriched in layers 4 and 6, whereas the influence of figure-ground segregation and visual attention is more pronounced in the superficial layers 2 and 3 and in layer 5 [27,60]. Thus, the feedforward group could correspond to neurons in layers 4 and 6 while the recurrent group could correspond to neurons in layer 2, 3 and 5. This view is supported by recent experimental evidence for the existence of separate channels for feedforward and feedback processing, suggesting that these influences might be mediated by partially non-overlapping neuronal circuits [61].

A study by Grossberg and Raizada [62] examined the propagation of enhanced activity along the representation of a target curve through horizontal connections in a model of the visual cortex. Their model also addressed the connectivity between cortical layers and predicted that the attentional effect would be relatively homogeneous across these layers, which does not match the observation that the attentional response enhancement is pronounced in the superficial and deep layers and weaker in layer 4 and 6 [27].

More recently, Marić & Domijan [63,64] configured a winner-take-all model to trace curves, also using separate feedforward and recurrent units. Their model could explain why tracing proceeds faster for target curves that are farther from a distractor curve than for target curves that are near a distractor, in accordance with psychophysical and neurophysiological data [18,65]. To propagate the enhanced activity, the model used a comparison between the activity levels of recurrent and feedforward units, whereas in the present model the activity propagation by recurrent units is gated by the feedforward units. This gating process prevents that recurrent activity spills over to units that represent the blank screen, helping with the selectivity of the propagation of activity among units representing connected image elements. The model of Marić & Domijan [63] also included units that were selective for the orientation of contours and it could therefore even trace target curves that crossed a distractor, based on the collinearity of contour elements. Orientation selectivity was lacking from the models studied by us, but future work could examine whether RELEARNN can train networks with orientation selective units to group collinear contour elements and disregard crossing distractor curves. It would also be interesting to test the spread of enhanced activity in artificial intelligence models with deeper feature hierarchies [66,67].

Linsley et al. [68] trained a recurrent neural network on a pathfinder task, which resembles curve-tracing, but now the curves consists of shorter, disconnected line elements. Their network had to determine whether two dots were on the same path. The authors demonstrated that a recurrent unit, called h-GRU [69], which activates based on feedforward connections but also integrates horizontal input, was useful to solve the task. Unlike the present study, the authors used backpropagation through time, which is a non-local learning rule that requires a teacher, unlike RELEARNN, which is a biologically plausible (also see [46]).

Whereas the models mentioned so far used enhanced activity to label image elements that are part of the same curve, Chen & Weng [70] modeled synchronous oscillations for perceptual grouping, in accordance with earlier theories on the role of oscillations in feature binding [71]. However, more recent neurophysiological studies demonstrated that synchrony and oscillations do not play a role in contour grouping [72], undermining their role in perceptual organization [73].

Our model extends previous modeling studies by demonstrating (1) the learning of visual routines based on curve-tracing and visual search and (2) how networks can be trained to execute visual routines using the biologically plausible RELEARNN rule. RELEARNN is equivalent to the Almeida-Pineda algorithm [36,37] and is local both in space and in time so that all the information that is necessary for the synaptic changes are available at the synapse. The learning rule relies on two networks. The first network is active during the processing of the stimulus when activity propagates through feedforward and recurrent connections. The second network is an accessory network, dedicated to help with the learning process. Activity in the accessory network is initiated by the action that is selected in the output layer and it then propagates through the network, emphasizing the synapses that have a strong influence on the activity of the winning output unit. Future neuroscientific work could test whether the proposed mechanisms are indeed implemented in the brain [74,75]. Firstly, the weights in the accessory network are proposed to be proportional to those of the main network. Previous work demonstrated that such symmetrical weights emerge during learning using the proposed learning scheme [76] and recent studies have suggested that the brain may even have specialized learning rules to promote this reciprocity [77]. In the brain, the approximate reciprocity of connections may hold at the level of cortical columns, but not at the level of individual neurons. Hence, the units in the simulated networks should be identified with cortical columns that consist of hundreds of cells and not with individual neurons.

In RELEARNN activity propagation occurs via regular recurrent units and the accessory units propagate a credit assignment signal. Both types of activity propagation are presumably related to visual attention at a psychological level of description. Some support for such different attentional networks comes from a study in area V4 of the visual cortex of monkeys by Steinmetz et al. [45], who used a task in which the animals had to visually search for a target item and had to make an eye movement to an opposite location in space. Some V4 neurons enhanced their activity when their RF was on the target of search, which would correspond to the regular recurrent units of the present study. Other V4 neurons enhanced their activity if their RF was on the target for the eye movement, which may correspond to the accessory units that play a role in movement selection and learning. The present work suggests that the propagation of enhanced activity by the regular units and the propagation of credit assignment signals by accessory units, in the opposite direction, might be separable functions of the attentional networks. We do realize, however, that the distinction between these two forms of attention is speculative and that much remains to be learned about the flow of attentional selection signals in brain circuits.

When the activity in the regular and accessory networks is combined with the reward prediction error, which can be mediated by a globally released neuromodulator such as dopamine, all the information for the synaptic update is available locally at the synapse. In previous work RELEARN was shown to train networks in the curve-tracing task [32] and here we showed that it is powerful enough to train networks on more complex, multistep tasks.

It is of interest that the networks developed strategies like those observed in the visual cortex of monkeys. During curve-tracing, recurrent units in the lower and higher layers propagated enhanced activity along the representation of the target curve, and it did not perform a sequence of discrete attention shifts, as was proposed by a previous model [31]. The gradual propagation of neuronal activity resembles how curve-tracing is implemented in the visual cortex of monkeys, where the neurons with RFs on the target curve successively enhance their activity, starting at the beginning of the curve until the end of the curve is reached. These responses enhancements are coordinated across multiple cortical areas including the frontal cortex, where the correct eye movement is selected [52]. Furthermore, the model implemented visual search by enhancing the activity of units coding for the relevant color, which then ultimately led to an enhanced representation of the location where this color was found, just as is observed in the visual cortex [19–24]. Finally, this propagation of enhanced activity was used by the model to transfer information from one elemental operation to the next, just like in the visual cortex of monkeys.

We here used a relatively simple search task for a specific color and we did not investigate whether the networks would exhibit increased processing delays in the presence of multiple color distractors or faster processing if the target is salient, which occurs in psychophysical experiments with human observers (e.g., [79]). Modeling accuracy and reaction times of human observers during visual search using network architectures like the one used here could be an interesting topic for future research.

Future modeling studies may also examine tasks in which the surfaces of 2D objects need to be grouped in perception. For example, Jeurissen et al. [80] found that the time it takes to determine whether two dots are on the same surface increases with the distance between the dots. The same principle that enables our network to trace curves could be expanded to enable networks to group 2D image regions, which are important in real-life scenarios, and the principles of region labeling might even be generalized to more complex situations that require semantic segmentation. Whereas we trained neural networks to carry out specific visual routines, it will be of interest to also explore scenarios in which elemental operations need to be organized in different sequences for a larger variety of tasks, e.g. considering the flexibility of humans who use a map to navigate. Future studies could also examine whether training on different tasks is possible if networks are deeper, so that the representations that select successive elemental operations that are part of a routine might emerge at higher network levels [3].

In conclusion, the new model architecture illustrates the versatility of networks that spread enhanced activity to solve elemental operations and to glue them together into visual routines, ensuring the transfer of the relevant parameters. The similarity between the network activity and neurons in the visual cortex indicate that the approach provides insight into how the circuits in the visual brain can configure themselves during learning, based on biological plausible learning rules that combine attentional and neuromodulatory signals. Future studies could expand this approach to further elucidate how feedforward, feedback and horizontal connections are used to enhance the cognitive capabilities of brain networks.

## Methods

### Data Analysis

We trained 24 networks for each task, which was the maximal number of networks that we could train in parallel. For the further analysis, we only kept networks that learned the tasks for curves up to 9 pixels long. We froze the weights after training and set the exploration rate *∈* to 0. We presented 15,000 randomly generated stimuli to each network and analyzed the activity of the units for these stimuli, including only correct trials in the analysis.

We placed the markers that related to the search tasks outside the main input grid, because incorporating them led to reduced stability. We note, however, that this stimulus design does not alter the structure of the tasks, or the need to transfer information between subroutines to correctly solve the search-then-trace and the trace-then-search tasks.

### Parameters

All networks were initialized and trained with the same parameters. They are summarized in the following table

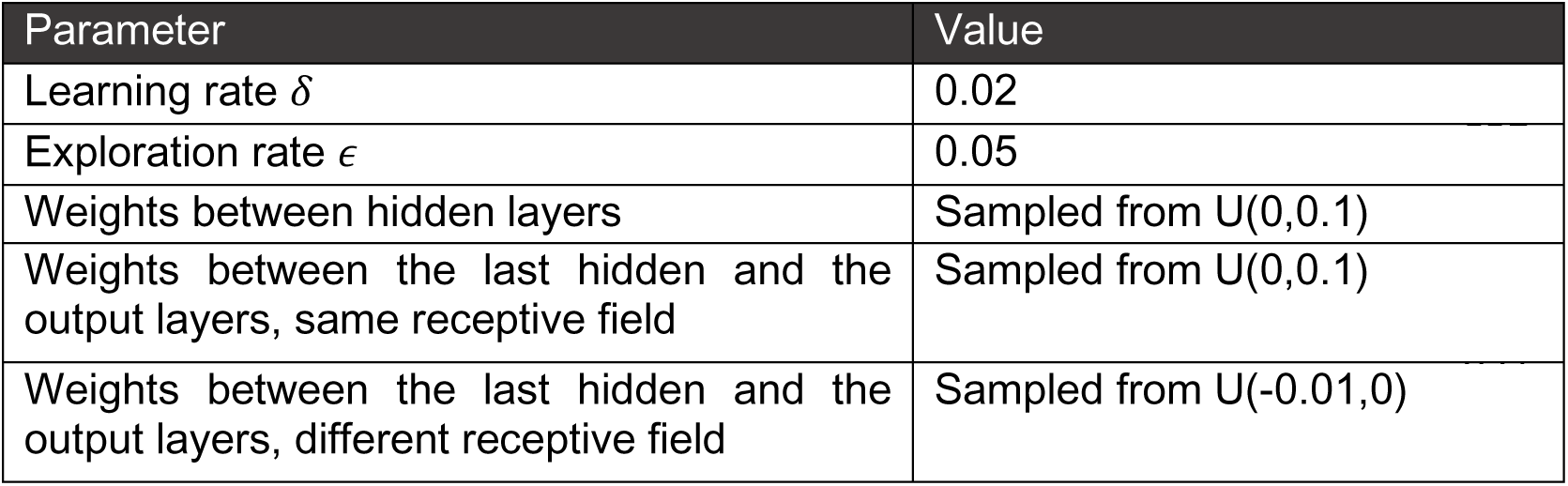

| Parameter | Value |
| --- | --- |
| Learning rate $\delta$ | 0.02 |
| Exploration rate $\epsilon$ | 0.05 |
| Weights between hidden layers | Sampled from $U(0,0.1)$ |
| Weights between the last hidden and the output layers, same receptive field | Sampled from $U(0,0.1)$ |
| Weights between the last hidden and the output layers, different receptive field | Sampled from $U(-0.01,0)$ |

Here U(x,y) refers to the uniform distribution on the interval [x,y].

### Code availability

All the code used to train the networks and to analyze the data is available on the following GitHub address: https://github.com/samimol/Visual-Routines

## Acknowledgements

This research has received funding from the European Union’s Horizon 2020 Framework Programme for Research and Innovation under the Specific Grant Agreement No. 945539 (Human Brain Project SGA3, Task 3.7), an ERC grant (101052963 “NUMEROUS”), an NWO Crossover grant 17619 “INTENSE”, “DBI2” (a Gravitation program of the Dutch Ministry of Science) and NWO-OCENW.KLEIN.178.

We acknowledge the use of Fenix Infrastructure resources, which are partially funded from the European Union’s Horizon 2020 research and innovation programme through the ICEI project under the grant agreement No. 800858.

---

**Fig. S1.**
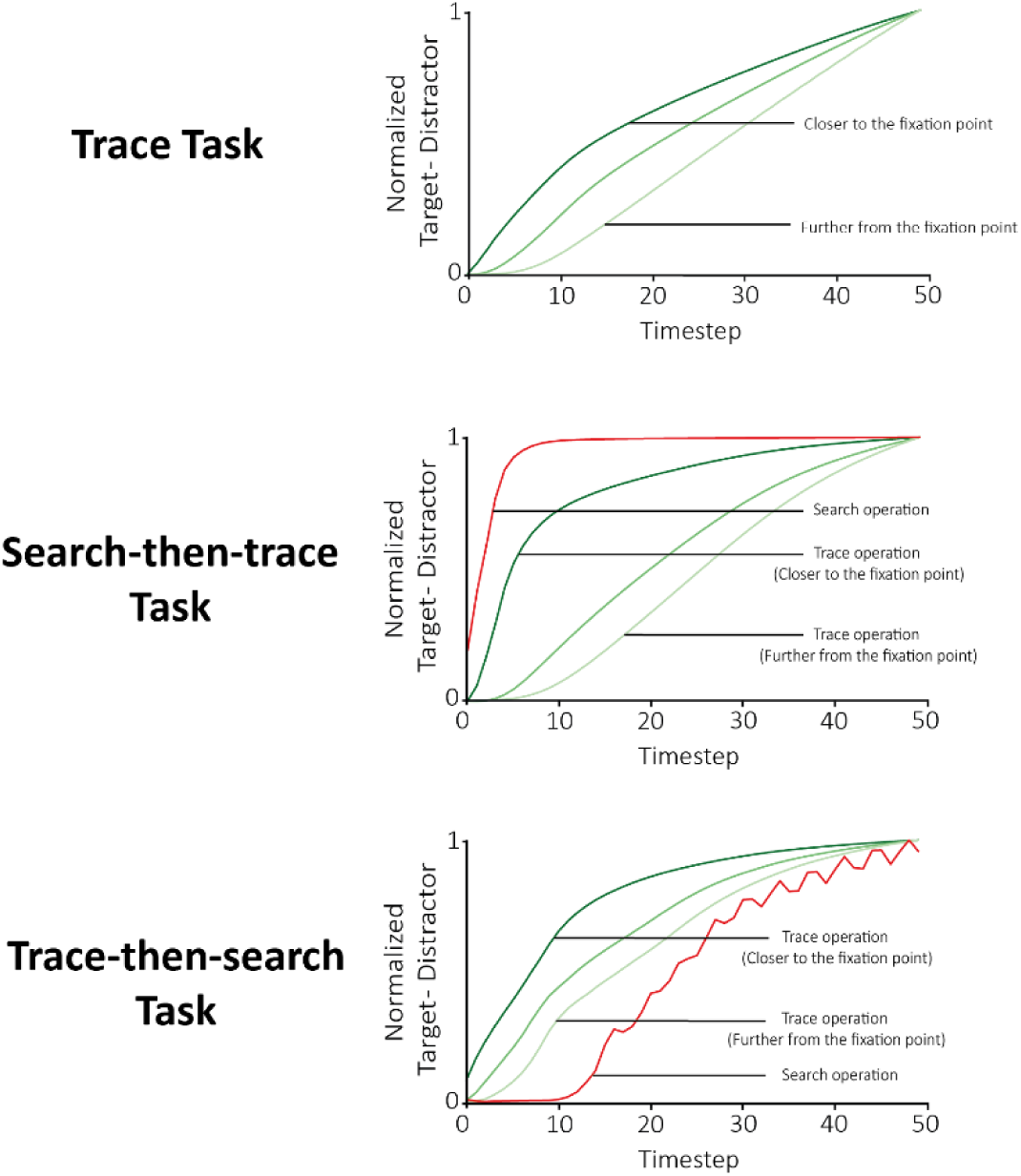
Time course of activity modulation in the output layer. In the output layer, the enhancement of activity of units with RFs on the target curve followed the same time course as neurons in the lower layers. Hence, the time required for spreading activity in the visual layers predicts the reaction time. These results are compatible with the reaction time measurements in humans by Jolicoeur et al. (1986). They demonstrated that the reaction time increases in proportion to the length of the curve that needs to be traced.

